# A non-destructive approach to assess the gut microbiome of honey bee (*Apis mellifera*) queens using fecal samples

**DOI:** 10.1101/2025.03.26.645508

**Authors:** Erick V. S. Motta, Jane Seong, Mustafa Bozkus, Tonya F. Shepherd, Juliana Rangel

## Abstract

Fecal sampling is a widely used, non-invasive method for assessing gut microbiomes across various organisms. However, its suitability for studying the gut microbiome of honey bee (*Apis mellifera*) queens has not been tested. In this study, we evaluated whether fecal microbiomes accurately reflect gut microbiomes in honey bee queens, offering a potential non-destructive approach for microbiome research. We successfully obtained fecal and gut samples from 21 out of 26 queens. Bacterial communities were analyzed using 16S rRNA amplicon sequencing and qPCR. Our results indicate that queen fecal microbiomes closely resemble gut microbiomes, with no significant differences in alpha diversity and only minor differences in specific bacterial taxa. Beta diversity analyses revealed that within-pair microbiomes (i.e., gut vs. feces from the same queen) were significantly more similar than between-pair comparisons. Additionally, qPCR analyses revealed a strong positive correlation between bacterial abundances in fecal and gut samples, further supporting the use of feces as a proxy for gut microbiome composition. While promising, fecal collection from queens can sometimes be challenging. In our study, we were unable to collect feces from five queens, and these individuals lacked stored fecal material upon inspection of dissected guts. Nonetheless, our findings suggest that fecal sampling can be a useful, non-invasive method for studying honey bee queen microbiomes, enabling longitudinal assessments without compromising colony stability.

**Importance:** The gut microbiome plays important roles in honey bee health, including protection against pathogens and stimulation of host physiological pathways. Because gut microbiome analysis involves destructive dissections, most research has focused on workers, given that colonies can have tens of thousands of individuals. This has left a significant gap in microbiome studies of other colony members, such as queens. Given that a queen is the sole reproductive female, her gut microbiome cannot be sampled without compromising colony integrity. In this study, we demonstrate that fecal sampling from honey bee queens provides a reliable, non-destructive alternative for assessing the composition of their gut microbiomes. We found that queen fecal microbiomes closely resemble their gut microbiomes in both taxonomic diversity and composition. This approach enables longitudinal monitoring of queen microbiomes without disrupting colony integrity, offering a valuable tool for honey bee research.

## Introduction

Fecal microbiomes are commonly used to study gut microbiomes due to their ease of collection and non-invasive nature (1). Fecal samples often provide a good representation of gut microbial communities, particularly those from the distal gut, making them a practical choice for microbiome research. Their collection is logistically simple, cost effective, and allows for repeated sampling, which is especially beneficial for longitudinal studies (2). However, fecal samples may not fully capture microbial diversity, particularly for microbes residing in the proximal gut or those attached to mucosal layers (1, 3). Therefore, direct comparisons between fecal and gut microbiomes should be made whenever possible to better assess the extent to which fecal samples accurately represent gut microbial communities.

In honey bee (*Apis mellifera*) research, microbiome studies typically involve dissecting the gut to obtain DNA for metagenomic analysis, providing a detailed view of an individual’s microbial composition, including specific gut compartments (4–8). While this method is feasible for studies of worker and drone bees, given that a colony is comprised of thousands of these individuals (9), it poses a significant challenge for queens, whose removal can threaten colony stability due to their role as the sole reproductive female in the colony. As a result, queen microbiome studies are often limited to periods of queen rearing or cases where colonies (or their queens) are no longer needed and can be sacrificed (5, 10).

Previous studies have indicated that queen gutmicrobiomes are distinct from those of workers and drones (5, 10, 11), highlighting the need for targeted research. In honey bee workers, the gut microbiome is dominated by five socially acquired core bacterial genera: *Bifidobacterium*, *Bombilactobacillus*, *Gilliamella*, *Lactobacillus*, and *Snodgrassella*. These bacteria are transmitted through social interactions and remain consistent across age (12). This microbiota contributes to digestion, development, and pathogen protection (13). In contrast, the queen’s gut microbiome appears to be more influenced by environmental factors and consists primarily of *Apilactobacillus*, *Bombella*, *Commensalibacter*, and *Lactobacillus*, many of which are also found in nectar, larvae, and hive compartments (5, 10, 11, 14). While some of these bacteria, such as *Bombella*, are known to protect larvae against pathogenic fungi (15) and enhance nutrition through lysine supplementation (16), their specific roles in queen health remain poorly understood. Understanding the queen’s microbiome is essential, as it may influence her health and, consequently, the health of the entire colony.

Recent studies have explored the use of fecal material for queen genotyping (17) and worker gut microbiome analysis (18), providing a non-invasive alternative for longitudinal studies. However, its applicability for assessing the queen’s gut microbiome remains untested. Since fecal microbiomes may not fully reflect gut composition, comparative studies are necessary before implementing this method in large-scale experiments. Here, we tested whether queen fecal sampling can serve as a non-destructive approach to investigate the queen’s gut microbiome. We collected fecal samples and dissected the guts of honey bee queens to compare their microbiomes using high-throughput sequencing and qPCR. This approach allowed us to determine whether fecal microbiome profiles reliably represent gut microbiomes, potentially providing a valuable non-destructive assessment method for honey bee queen microbiomes.

## Results

### Collection of fecal samples from honey bee queens

To determine whether honey bee queen feces harbor bacterial communities similar to those found in their guts, we collected fecal samples before gut dissections (Figure 1). Collection was easy for some queens, but challenging for others. Of the 26 queens sampled, we obtained at least 1 µL of feces from 21 queens (81% of the total samples; Table S1). We then dissected the guts of the queens from which we successfully collected feces, yielding 21 paired feces/gut samples, which were processed alongside three worker gut samples and a negative control, which contained all components of the extraction protocol except for the gut or feces, for 16S rRNA amplicon sequencing and qPCR analysis.

**Figure 1.**
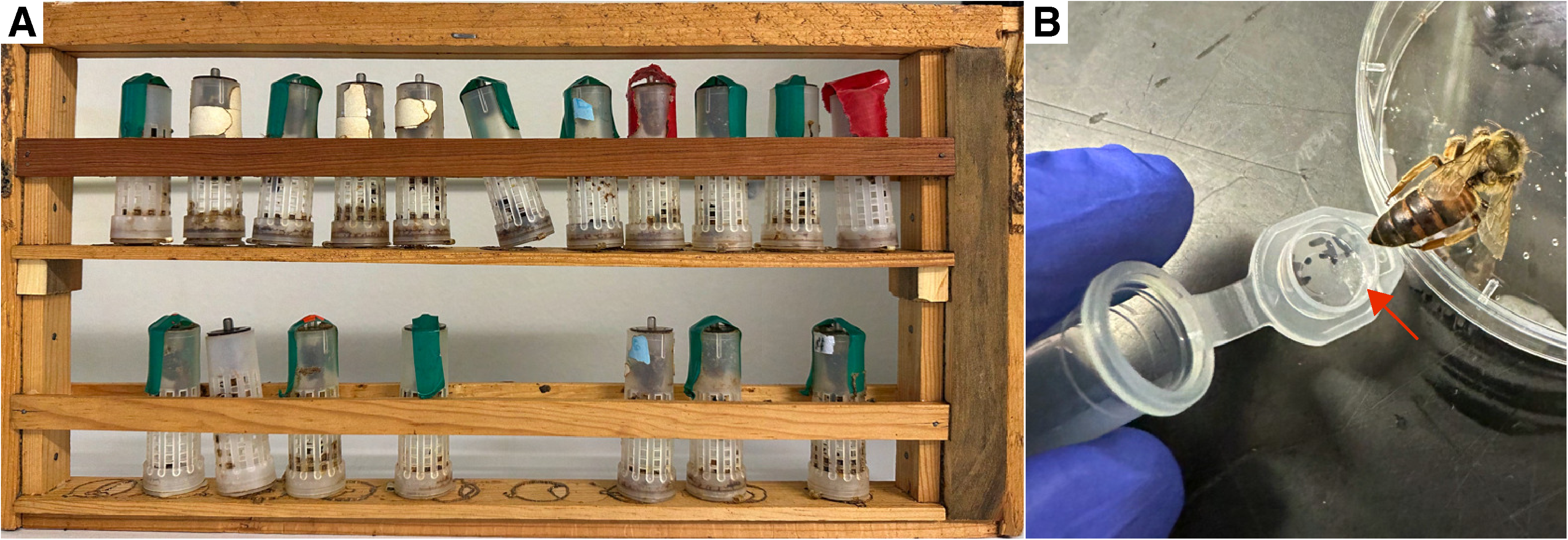
Honey bee queen grafting and feces sampling. **(A)** Individual queens were raised in plastic cups via grafting. Once the cells were capped, they were individually placed in plastic “roller” cages and placed in a queen “bank” where they were taken care of by tending bees until collection for analysis. **(B)** Feces were sampled from chilled queens by positioning the distal region of the abdomen toward the cap of a sterile microcentrifuge tube and gently pressing the abdomen with the investigator’s index and thumb fingers until feces were released. The queen feces were mostly clear, as indicated by the red arrow.

### Bacterial diversity in fecal and gut samples

Based on 16S rRNA amplicon sequencing, we found that bacterial communities in queen feces were similar to those detected in their respective guts (Figure 2A, Figure S1). The main bacterial genera detected in both types of samples included species of *Apilactobacillus*, *Bifidobacterium*, *Bombella*, *Bombilactobacillus*, *Commensalibacter*, and *Lactobacillus*, which is consistent with findings from other studies on queen gut microbiota (5, 10, 11).

**Figure 2.**
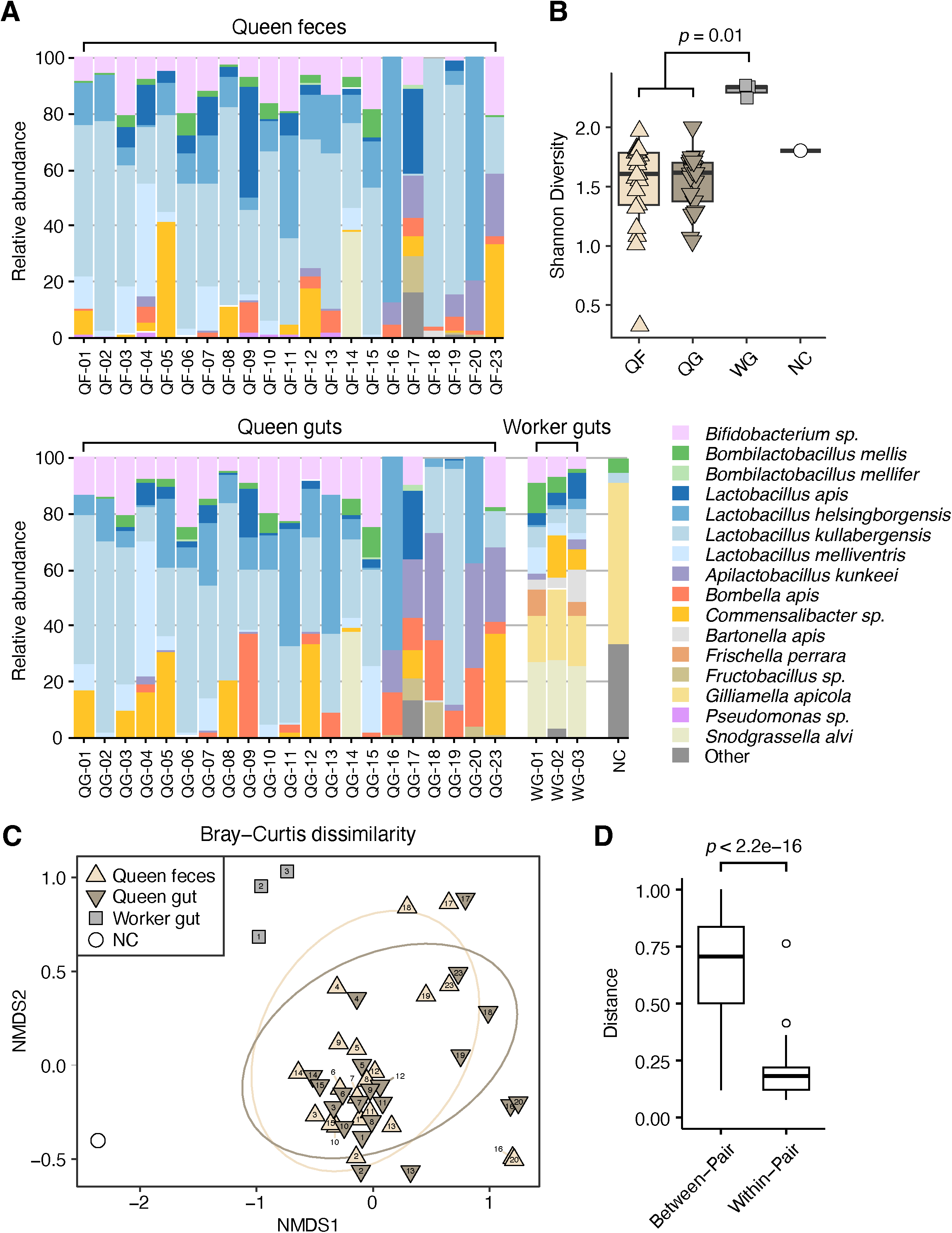
Comparisons of the relative abundance and microbial diversity of honey bee queen fecal and gut microbiomes. **(A)** Stacked column graphs showing the relative abundance of gut bacterial species in queen feces (QF, n = 21), queen guts (QG, n = 21), worker guts (WG, n = 3), and a negative control (NC, n = 1) to check for contamination during library preparation. Fecal and gut samples with the same number originated from the same queen. **(B)** Shannon diversity index comparing alpha diversity across groups. Statistical analysis was performed using the Kruskal-Wallis test (χ^2^ = 8.58, df = 2, *p*-value = 0.01, n = 45), followed by Dunn’s pairwise comparisons with *p*-values adjusted using the Benjamini-Hochberg method (*p*-value adj < 0.05 shown). NC was excluded from the analysis. **(C)** Non-Metric Multidimensional Scaling (NMDS) based on Bray-Curtis dissimilarity of gut community compositions. Statistical analysis was performed using PERMANOVA (F_3,42_ = 3.52, permutations = 999, *p*-value = 0.01), followed by pairwise comparisons. Significant differences were observed between QF vs. WG (*p*-value adj = 0.012) and QG vs. WG (*p*-value adj = 0.006). **(D)** Pairwise distance analysis using Bray-Curtis dissimilarity to compare within-pair (gut vs. feces from the same queen) and between-pair (samples from different queens) comparisons (*t*-test, *p*-value < 2.2e-16).

Comparisons of microbial diversity (both alpha and beta) between queen feces and guts demonstrated a high degree of similarity. For alpha diversity, which measures diversity within samples, we examined the Shannon diversity index (accounting for both species richness and evenness) and found a significant difference among the groups (Kruskal-Wallis test, χ^2^ = 8.58, df = 2, N = 45, *p*-value = 0.01). Post hoc analysis indicated that both queen feces and queen gut samples exhibited significantly lower Shannon index compared to worker gut samples (Dunn’s test, Queen Feces vs. Worker Gut: Z = -2.63, *p*-value adj = 0.01; Queen Gut vs. Worker Gut: Z = -2.93, *p*-value adj = 0.01, Figure 2B); however, there was no significant difference between queen feces and gut bacterial communities (Dunn’s test, Z = 0.59, *p*-value adj = 0.55, Figure 2B). Additionally, a differential abundance analysis between queen gut and fecal microbiomes revealed that only two amplicon sequence variants (ASVs) belonging to the genera *Pseudomonas* and *Lactobacillus* were significantly enriched in queen feces compared to queen guts (DESeq2: *p*-value adj < 0.05, fold change > |2|, n = 42, Figure S2).

Regarding beta diversity, which measures diversity between samples, we used two metrics: Bray-Curtis dissimilarity (measuring differences in species abundances) and Weighted UniFrac (accounting for species abundances and their evolutionary relationships) (19, 20). NMDS plots based on these metrics were used for data visualization (Figure 2C and Figure S3A). PERMANOVA analyses indicated significant differences between groups for both metrics (Bray-Curtis: F_3,42_ = 3.52, permutations = 999, *p*-value = 0.01; Weighted UniFrac: F_3,42_ = 13.65, permutations = 999, *p*-value = 0.001). Pairwise comparisons revealed that these differences were primarily driven by comparisons between worker gut samples and queen samples, for both gut and feces (*p*-value adj < 0.05).

To investigate whether gut and fecal samples from the same individual were more similar compared to those from different individuals, we performed an additional pairwise distance analysis using Bray-Curtis and Weighted Unifrac distances. The analyses included both within-pair comparisons (gut vs. feces from the same individual) and between-pair comparisons (gut vs. feces from different individuals). Both the Bray-Curtis and Weighted Unifrac distances between gut and fecal samples from the same individual were significantly smaller than the distances between samples from different individuals, suggesting that gut and fecal microbiomes within the same individual (within-pair comparisons) were more similar to each other compared to those from different individuals (between-pair comparisons) (t-test, *p*-value < 2.2e-16, Figure 2D, Figure S3B).

### Bacterial loads in fecal and gut samples

When we quantified bacterial loads in our samples, we found that queen feces exhibited average bacterial loads of 1.08 × 10^8^ 16S rRNA amplicon copies per μL (95% CI: 4.07 × 10^7^ to 1.76 × 10^8^, N = 21), while queen gut samples exhibited average bacterial loads of 1.16 × 10^10^ 16S rRNA amplicon copies per gut (95% CI: 6.24 × 10^9^ to 1.69 × 10^10^, N = 21). Compared to worker gut samples, whose average bacterial loads were 2.86 × 10^9^ 16S rRNA amplicon copies per gut (95% CI: 6.79 × 10^8^ to 5.04 × 10^9^, N = 3), our findings also suggest that queen guts can harbor similar bacterial loads than worker guts (Figure 3A). A negative control sample indicated almost no external contamination of our processed samples (17.27 16S rRNA amplicon copies per μL).

**Figure 3.**
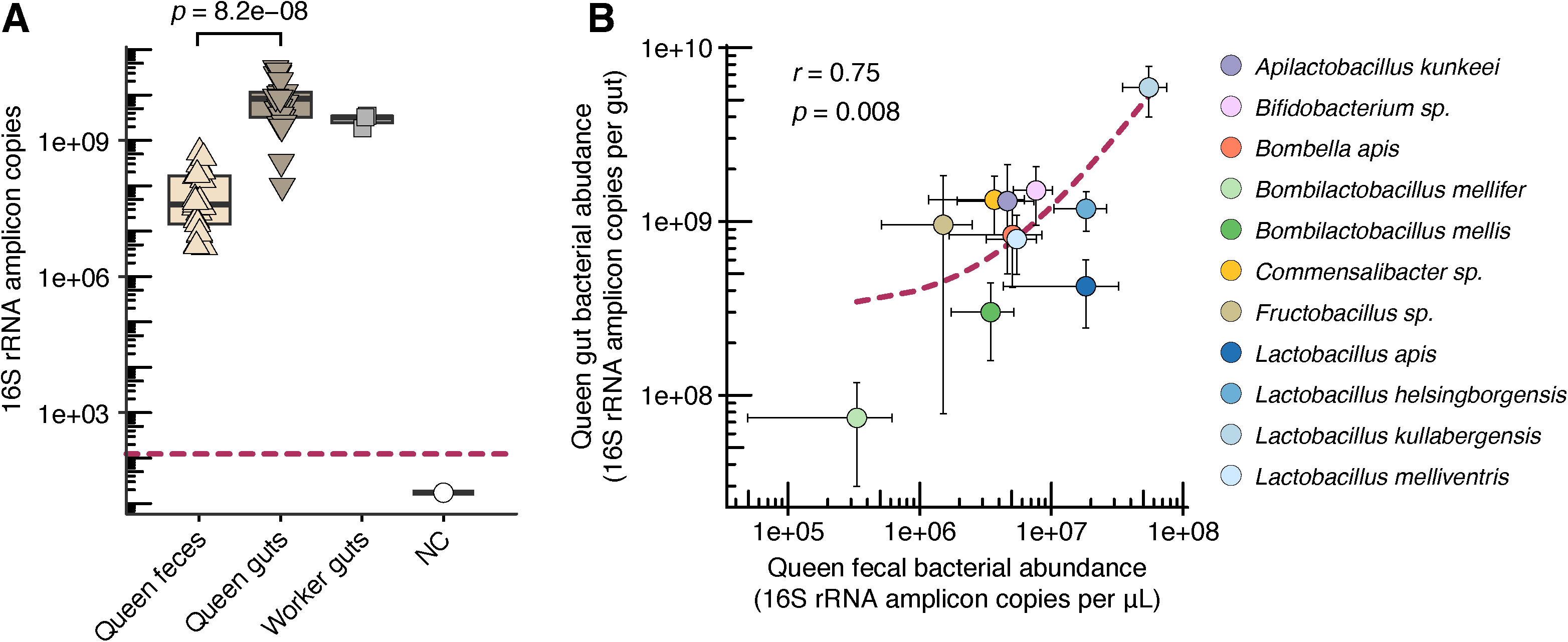
Comparisons of bacterial loads in honey bee queen fecal and gut microbiomes. **(A)** Box plots showing estimates of total bacterial abundance in queen feces (QF, n = 21), queen guts (QG, n = 21), worker guts (WG, n = 3), and a negative control (NC, n = 1). Statistical analysis was performed using Kruskal-Wallis test (χ^2^ = 31.23, df = 2, *p*-value = 1.65e-07, n = 45), followed by Dunn’s pairwise comparisons with *p*-values adjusted using the Benjamini-Hochberg method (*p*-value adj < 0.05 shown). The maroon dashed line indicates the limit of detection of the qPCR analysis. **(B)** Scatter plot showing the correlation between the absolute abundances of bacterial genera in feces and gut samples (Pearson correlation coefficient *r* and *p*-value shown). Only ASVs with absolute abundance above 0.1% in at least three samples are displayed for clarity. The maroon dashed line represents the linear regression curve, which appears as non-linear due to log axes.

Finally, when we combined information from the qPCR analysis with the 16S rRNA amplicon sequencing data, we found a positive correlation between the absolute abundance of the most common bacterial taxa found in queen guts and feces, indicating that the level of gut colonization by a specific taxon was mirrored by its abundance in the feces (Pearson correlation coefficient *r* = 0.75, N = 42, *p*-value = 0.008, Figure 3B, Figure S4). Overall, our findings suggest that fecal samples can be used for the study of queen gut microbiota composition, as they enabled us to determine the presence or absence of bacteria (community membership), as well as estimate bacterial loads (colonization levels) in the gut of individual queens in a non-destructive manner.

## Discussion

Our study demonstrates that fecal samples can serve as a reliable non-destructive method for assessing the gut microbiomes of honey bee queens. We found that bacterial communities in queen feces closely resemble those in their guts, with no significant differences in alpha or beta diversity and only minor differences in specific bacterial taxa. This similarity suggests that fecal samples can provide a representative snapshot of a queen’s gut microbiome, enabling longitudinal studies without harming the queens, similarly to what has been proposed for workers (18, 21). Additionally, we found a positive correlation between the absolute abundance of bacterial taxa in the feces and guts of queens, further supporting the use of fecal samples for queen microbiome analysis. This positive correlation has also been observed in a study comparing the bacterial loads in the guts and feces of honey bee workers (18).

Other studies, particularly in insects, have shown that fecal microbiomes can reflect gut microbiomes. For example, research on German cockroaches (22), silkworms (23), and spider mites (24) found significant overlap between microbial communities in the gut and feces, suggesting that fecal samples can provide valuable insights into insect gut microbiomes. However, this is not always the case, as studies on mammals indicate that fecal microbiomes may not accurately represent the gut microbiome of specific gut regions. A study on pigs analyzing microbial and metabolic compositions at multiple gastrointestinal sites found significant differences in microbiome diversity and metabolite composition (3), highlighting the need to compare fecal and whole-gut communities when assessing an organism’s microbiome.

The fecal sampling process we used in our study presented some challenges. While we successfully collected feces from most of the queens, the volume we obtained varied considerably (from one microliter to several microliters), and for some queens, no feces could be collected, even after applying considerable pressure on their abdomens. This variability in feces collection may be influenced by factors such as hydration, feeding status, or prior defecation while in cages, and brings an extra level of complexity to consider when designing experiments to assess queen fecal microbiomes. Moreover, our queens remained banked until processed and were never released to the hive. Therefore, it is possible that queens in a regular hive environment may have different levels of fecal material to be collected for these types of studies.

In our study, we immobilized queens by placing them inside clean, sterile Petri dishes and then on ice for several minutes until they were immobilized and processed for feces collection. Bubnič et al. (2020) (17) anesthetized queens using CO_2_ gas instead of cold temperature, then transferred them to a clean, sterile Petri dish, where the queens defecated naturally within minutes after waking up. On the other hand, Cabirol et al. (2024) (18) stunned workers (not queens) using CO_2_ and immobilized them on ice to collect feces and guts. In our study, we directly immobilized queens on ice. For the queens from which we could not obtain feces after strongly pressing their abdomens, we allowed them to return to room temperature and stay in the Petri dish for several minutes. However, this did not induce defecation later on. We then chilled them on ice again and dissected their guts for inspection, observing empty rectum sections. This suggests that even if we had exposed the queens to CO_2_, they would not have defecated as there were no feces stored in their rectums. Therefore, feeding queens prior to sampling could increase the chances of obtaining fecal material in specific cases.

In conclusion, fecal collection is a promising, non-destructive approach for the study of honey bee queen gut microbiomes. This method can facilitate longitudinal research to assess the impacts of environmental stressors, pathogens, diet, and other factors on queen health. Future studies should investigate the long-term effects of fecal induction and collection on queen survival and colony stability.

## Material and Methods

### Honey bee queen rearing

The colonies used in this study were maintained at the Janice and John G. Thomas Honey Bee Facility of Texas A&M University in Bryan, TX (N 30°38’31.037” W 96°27’ 39.495’”). All colonies were headed by queens of Italian descent. Experimental queens were reared by using the standard queen-rearing procedure known as “grafting” (25, 26), which entailed selecting a frame of developing brood, and transferring first instar worker larvae from their cells into plastic cups using a grafting tool (JZ’s BZ’s Honey Co., Santa Cruz, CA). We prepared the plastic cell cups by priming them with a small amount of royal jelly to encourage larval acceptance and grafted a total of 30 larvae per each of two grafting sessions, as shown in Figure 1. The grafted cell cups were then attached to a queen-rearing frame and placed in a queenless unit of bees known as a “cell builder” (25), where nurse bees cared for queens during larval and pupal development. Larvae were grafted during two different sessions, one on 7 July 2024 (first session) and the other on 11 July 2024 (second session). All emerged queens were placed in a “bank”, where they were taken care of by tending workers until they were used. Banked queens were brought to the laboratory for analysis on two different dates: eight queens from the second grafting session (approximately 38 days old) were analyzed on 29 August 2024, and 18 queens (13 from the first grafting session and five from the second grafting session, approximately 80-85 days old) were analyzed on 8 October 2024.

### Feces sampling and gut dissections

We used 26 queens for fecal sample collection. Each queen was individually immobilized in a clean, sterile Petri dish on ice for about 10 minutes. Then, queens were carefully handled by directing the postal region of the abdomen toward the cap of a sterile 1.5 mL microcentrifuge tube, after which light pressure was applied to their abdomen with the researcher’s index and thumb fingers (Figure 1). The total volume of feces collected from queens varied considerably, ranging from 0 µL to 10 µL. For the queens from which we did not obtain feces (N = 5), a stronger pressure was applied to the abdomen, which may have caused damage to the queen. As all these queens were going to be dissected for gut extractions, we were not worried about such damage, but if these queens were to be transferred back to a hive, this strong pressure should be avoided. For the queens from which we were able to collect at least 1 µL of feces (N = 21), we proceeded with the gut dissections. Guts were removed by using sterilized forceps by grabbing the last segment of the abdomen, allowing us to pull out the entire gut. Each gut was transferred to an individual sterile 1.5 mL microcentrifuge tube. All feces and gut samples were obtained under sterile conditions and immediately stored at -80 °C until further processing.

### DNA extraction

DNA was extracted from queen feces (n = 21), queen guts (n = 21), and worker guts (n=3) using a previously described protocol (7), with some adaptations. Briefly, guts were homogenized with 100 μL of CTAB buffer (0.1 M Tris-HCl pH 8.0, 1.4 M NaCl, 20 mM EDTA, and 2% (w/v) cetrimonium bromide), resuspended in additional 600 μL of CTAB buffer and 20 μL of proteinase K solution (0.1 M tris-HCl, 26 mM CaCl_2_, 50% glycerol, and 20 mg/mL proteinase K), and transferred to a capped vial with 0.5 mL of 0.1 mm Zirconia beads (BioSpec Products). After adding 2 μL of 2-mercaptoethanol, samples were bead-beated for 2 × 2 min. Samples were digested at 56 °C overnight, transferred to a clean 2 mL vial and mixed with 500 μL of phenol-chloroform-isoamyl alcohol (25:24:1, pH 8.0). Samples were inverted five times and centrifuged for 15 min at 4 °C and 18,000 RCF (Eppendorf). The aqueous layer was transferred to a new vial, and DNA was precipitated at −20 °C overnight with 500 μL of isopropanol and 50 μL of 3 M NaOAc (pH 5.4). Precipitated samples were centrifuged for 30 min at 4 °C and 18,000 RCF (Eppendorf), and the supernatant was removed. DNA pellets were washed with 1 mL of ice cold 75% ethanol and centrifuged for additional 3 min at 4 °C. After removing the ethanol wash, the DNA pellets were dried at room temperature for 30 min and then resuspended in 50 μL of water. Final DNA samples were stored at −20 °C.

### qPCR analysis

The DNA samples from feces were used directly as templates for qPCR, while gut DNA samples were diluted 10-fold. Triplicate reactions were prepared in 96-well plates (one for feces, one for gut samples) and run on a Bio-Rad CFX96 Touch Real-Time PCR instrument. Each reaction contained 5 μL SYBR Green Universal Master Mix (Applied Biosystems), 0.05 μL of 100 μM forward and reverse primers (27F: 5’- agagtttgatcctggctcag-3’ and 355R: 5’-ctgctgcctcccgtaggagt-3’), 3.9 μL H_2_O, and 1.0 μL template DNA. Cycling conditions included an initial step at 50 °C for 2 min and 95 °C for 2 min, followed by 40 cycles of 95 °C for 15 s and 60 °C for 1 min.

Total bacterial 16S rRNA gene copies were quantified using standard curves generated from a synthesized 347-bp 16S rRNA gene fragment (Twist Bioscience). The fragment was quantified (Qubit) and adjusted to 10^10^ copies/μL. Serial dilutions from 10^8^ to 10^2^ copies/μL were prepared as standards and used in each plate. 16S rRNA gene copy numbers were calculated as 10^(Ct-b)/m^, where “b” and “m” are the y-intercept and slope of the standard curve, respectively, and Ct (cycle threshold) was the average of triplicates. Values were corrected for the dilution factor.

### 16S rRNA amplicon sequencing

DNA samples from feces and 10-fold diluted DNA samples from guts were used as templates for 16S rRNA library preparation, involving two PCR reactions. PCR 1 amplified the V4 region of the 16S rRNA gene in 25 μL single reactions, including 1 μL of 10 μM forward and reverse primers (Hyb515F: 5’- tcgtcggcagcgtcagatgtgtataagagacaggtgycagcmgccgcggta-3’ and Hyb806R: 5’- gtctcgtgggctcggagatgtgtataagagacagggactachvgggtwtctaat-3’), 12.5 μL of 2× AccuStart^TM^ II PCR SuperMix (Quantabio), and 1 μL of template DNA. Cycling conditions were 94 °C for 3 min; 30 cycles of 94 °C for 20 s, 50 °C for 15 s, 72 °C for 30 s; followed by 72 °C for 10 min. PCR 2 attached dual indices and Illumina sequencing adapters to PCR 1 products in 25 μL single reactions, including a unique combination of 2 μL of 5 μM index primers (Hyb-F*nn*-i5: 5’- aatgatacggcgaccaccgagatctacacnnnnnntcgtcggcagcgtc-3’, and Hyb-R*nn*-i7: 5’- caagcagaagacggcatacgagatnnnnnngtctcgtgggctcgg-3’), 12.5 μL of 2× AccuStart^TM^ II PCR SuperMix, and 5 μL of PCR 1 product. Cycling conditions were 94 °C for 3 min; 10 cycles of 94 °C for 20 s, 50 °C for 15 s, 72 °C for 60 s; followed by 72 °C for 10 min.

Both PCR product sets were purified using 0.8× HighPrep PCR magnetic beads (MagBio) and quantified with a Qubit Fluorometer (Thermo Fisher Scientific). Samples (200 ng each) were pooled and submitted for Illumina sequencing on the MiSeq platform (2 × 250 bp run) at Texas A&M University’s Genomics & Bioinformatics Service (College Station, TX).

During library preparation, we included one negative control (N = 1) to check for contamination, which consisted of adding molecular biology grade water instead of a DNA sample for the first PCR reaction.

### 16S rRNA amplicon analysis

Illumina sequence reads were demultiplexed based on barcode sequences using MiSeq software. Read quality was assessed using FastQC (27) and MultiQC (28). Then, reads were processed in QIIME 2 version 2024.5 (29) on the Grace cluster at the Texas A&M High-Performance Research Computing (HPRC) facility. Paired-end FASTQ files were imported into QIIME 2 using the Casava 1.8 paired-end demultiplexed FASTQ format. Primer sequences (V4 region: gtgycagcmgccgcggta and ggactachvgggtwtctaat) were removed using the cutadapt plugin (30). Reads were filtered for quality, truncated to 220 bp, denoised, merged, and chimeric reads were removed, all using the DADA2 plugin. (31).

Taxonomy was assigned to amplicon sequence variants (ASVs) using the SILVA 138 reference database in the feature-classifier plugin (32). To validate taxonomic assignments when needed, representative sequences were examined using BLASTn against the NCBI database. Low-abundance reads (<1% relative abundance), unassigned sequences, and non-bacterial reads (mitochondrial and chloroplast) were filtered using the feature-table and taxa filter-table plugins. An ASV table was generated to investigate microbial composition across all samples and exported for data visualization and statistical analysis in R version 4.3.2 (33).

### Statistical analysis

Microbial diversity analyses were performed using the R package phyloseq (34). An ASV table, phylogenetic tree, and mapping file were imported and merged into a single phyloseq object. Taxonomy rank columns were assigned, and read counts were scaled to an even depth using a custom function (35). For alpha diversity, Shannon index was calculated and differences between groups were assessed by performing a Kruskal-Wallis test followed by Dunn’s test using the R package FSA (36). For beta diversity, ordinations and distance matrices were calculated using Bray-Curtis and Weighted UniFrac metrics and visualized as Non-Metric Multidimensional Scaling (NMDS) plots. PERMANOVA was conducted to analyze differences between groups using the adonis2 function from the R package vegan (37), and pairwise comparisons were performed with pairwise.adonis function from the same package. To investigate whether gut and fecal samples from the same individual (or pair) are more similar than those from different individuals, we extracted within-pair and between-pair distances from both beta diversity metrics using the R package phyloseq (34), then performed t-tests using the R package ggpubr (38).

For differential expression analysis, we used the R package DESeq2 (39). The phyloseq object was converted to a DESeq2 object, and size factors were estimated to normalize for sequencing depth. We ran the DESeq2 analysis to test for differential abundance between groups and extracted significant ASVs. Results were visualized using R package ggplot2 (40), creating a volcano plot and boxplots for significant ASVs. We set the level of statistical significance for all tests at α = 0.05.

## Supplementary figure legends

**Figure S1.** Boxplots showing relative abundances of representative bacterial genera in queen feces (n = 21), queen guts (n = 21), worker guts (n = 3), and a negative control (NC) based on 16S rRNA amplicon data. Statistical analysis was performed using Kruskal-Wallis tests followed by Dunn’s pairwise comparison tests, with *p*-values adjusted using the Benjamini-Hochberg method (*p*-value adj < 0.05 shown).

**Figure S2.** Significant microbiome differences between queen feces and gut samples. **(A)** Volcano plot showing significance versus fold-change based on ASV relative abundances in queen guts compared to queen feces. Red dots indicate ASVs significantly different in DESeq2 analyses (FDR-corrected *p*-value < 0.05). **(B)** Boxplots depicting the relative abundance of significantly different ASVs between queen feces and gut samples, along with worker gut (WG) and negative control (NC) samples for comparison.

**Figure S3.** Non-Metric Multidimensional Scaling (NMDS) based on Weighted UniFrac dissimilarity of gut community compositions. Statistical analysis was performed using PERMANOVA (F_3,42_ = 13.65, permutations = 999, *p*-value = 0.001), followed by pairwise comparisons. Significant differences were observed between QF vs. WG (*p*-value adj = 0.006) and QG vs. WG (*p*-value adj = 0.006).

**Figure S4.** Boxplots showing absolute abundance estimates of representative bacterial genera in queen feces (n = 21), queen guts (n = 21), worker guts (n = 3), and a negative control (NC), based on qPCR. Statistical analysis was performed using Kruskal-Wallis tests followed by Dunn’s pairwise comparison tests, with *p*-values adjusted using the Benjamini-Hochberg method (*p*-value adj < 0.05 shown).

## Supplementary tables

**Table S1.** Approximate volume of feces collected from each experimental queen. “QF” stands for “Queen Feces”. Extraction batch #1 consisted of 13 queens from grafting session #1 and 5 queens from grafting session #2, and extraction batch #2 consisted of 8 queens from grafting session #2.

| Extraction batch #1 |  | Extraction batch #2 |  |
| --- | --- | --- | --- |
| Queen ID | Feces (μL) | Queen ID | Feces (μL) |
| QF-1 | 3.0 | QF-16 | 8.0 |
| QF-2 | 4.0 | QF-17 | 1.5 |
| QF-3 | 1.5 | QF-18 | 1.0 |
| QF-4 | 2.5 | QF-19 | 1.0 |
| QF-5 | 5.5 | QF-20 | 1.0 |
| QF-6 | 5.8 | QF-21 | - |
| QF-7 | 10.0 | QF-22 | - |
| QF-8 | 8.5 | QF-23 | 5.0 |
| QF-9 | 2.0 |  |  |
| QF-10 | 1.0 |  |  |
| QF-11 | 4.5 |  |  |
| QF-12 | 2.8 |  |  |
| QF-13 | 3.5 |  |  |
| QF-14 | 1.8 |  |  |
| QF-15 | 1.5 |  |  |
| QF-16 | - |  |  |
| QF-17 | - |  |  |
| QF-18 | - |  |  |

## Data availability

16S rRNA amplicon sequencing data are available on NCBI BioProject PRJNA1234195. Other data are included in this article and its supplementary information files.

## Acknowledgements

We thank Keegan Nichols and Sam Gracia for helping us collect queens, and Jayda Arriaga for helping us check DNA quality. Portions of this research were conducted with the advanced computing resources provided by Texas A&M High Performance Research Computing.

